# GIVE: toward portable genome browsers for personal websites

**DOI:** 10.1101/177832

**Authors:** Xiaoyi Cao, Zhangming Yan, Qiuyang Wu, Alvin Zheng, Sheng Zhong

## Abstract

Growing popularity and diversity of genomic data demands portable and versatile genome browsers. Here, we present an open source programming library, called GIVE that facilitates creation of personalized genome browsers without requiring a system administrator. By inserting HTML tags, one can add to a personal webpage interactive visualization of multiple types of genomics data, including genome annotation, “linear” quantitative data (wiggle), and genome interaction data. GIVE includes a graphical interface called HUG (HTML Universal Generator) that automatically generates HTML code for displaying user chosen data, which can be copy-pasted into user’s personal website or saved and shared with collaborators. The simplicity of use was enabled by encapsulation of novel data communication and visualization technologies, including new data structures, a memory management method, and a double layer display method. GIVE is available at: http://www.givengine.org/.

## INTRODUCTION

Genomics data have become increasingly popular and diverse, posing new challenges to personalized data management and visualization [1-4]. On the one hand, people interested in making their genomic data public required “researchers and policy makers [to anticipate] when people share their genome on Facebook” [5]. This movement asks for development of portable, versatile, and easily deployable genome browsers. Ideally, a portable data visualization tool can work like Google map, that can be inserted into personal websites. On the other hand, new data types especially those representing genome-wide interactions, including genome-interaction data (Hi-C [6], ChIA-PET [7]), transcriptome-genome interaction data (MARGI [8], GRID-seq [9]) and transcriptome interaction data (PARIS [10], MARIO [11], LIGR-seq [12], SPLASH [13]) require compatible visualization tools, and ideally these data should be able to seamlessly displayed in parallel to other data types including RNA-seq [14], ChIP-seq [15], ATAC-seq [16].

It was envisioned that future genome browsers could work like Google map, of which users with small efforts can insert a customized version into their own websites [17]. Redeployable genome browsers are developed toward this goal [17-20]. Still, releasing websites with interactive visualization of genomic data would generally require systems administration, database and web programming work. The GIVE project is aimed to automate these work and offer a portable and lightweight genome browser with complementary advantages of genome browser websites [1, 21], desktop executables [22], and personal homepages and blogs.

We created the open source GIVE programming library to meet diverse needs of users with various levels of sophistication. A feature called GIVE HUG (<u>H</u>TML <u>U</u>niversal <u>G</u>enerator) provides a graphical interface to interactively generate HTML codes for displaying user chosen datasets. Users can save and share the HTML file with collaborators or copy-paste the HTML codes into her/his websites, which would lead to embedded interactive data display. Users can use GIVE to create custom genome browsers without hosting a data server, where all the data are retrieved on-demand from public data servers. Users who choose to host data on their own server can do so with commands provided in GIVE-Toolbox. With a few lines of HTML codes, GIVE enables a website to retrieve, integrate, and display diverse data types hosted by multiple servers, including large public depositories and custom-built servers. Such simplicity of use comes from encapsulation of new data management, communication, and visualization technologies made available by the GIVE development team. The cores of these technologies are new data structures and a memory management algorithm.

## RESULTS

### Overview of the GIVE library

GIVE is composed of an HTML tag library and GIVE-Toolbox. The former is a library of HTML tags for data visualization. GIVE-Toolbox is a set of command line commands, which automates all necessary database operations. For any public datasets for which the metadata can be found in GIVE data hub, users can directly use GIVE’s HTML tags to display such data, without invoking GIVE-Toolbox.

GIVE’s HTML tag library provides flexibility to build a variety of genome browsers, for example a single-cell transcriptome website [23] (https://singlecell.givengine.org/), an epigenome website [24, 25] (https://encode.givengine.org/, Figure S1), an genome interaction website [26] (https://mcf7.givengine.org/, Figure 1), and an RNA-chromatin interaction website [27] (https://margi.givengine.org/, Figure 2). With GIVE, users can build data visualization websites without hosting actual data (data are hosted on public data servers) or data hosting websites or websites that displays composite datasets hosted on user’s server and public servers. The GIVE-enabled HTML files can also be used and shared as custom software, which encapsulate both data and visualization capability.

**Figure 1.**
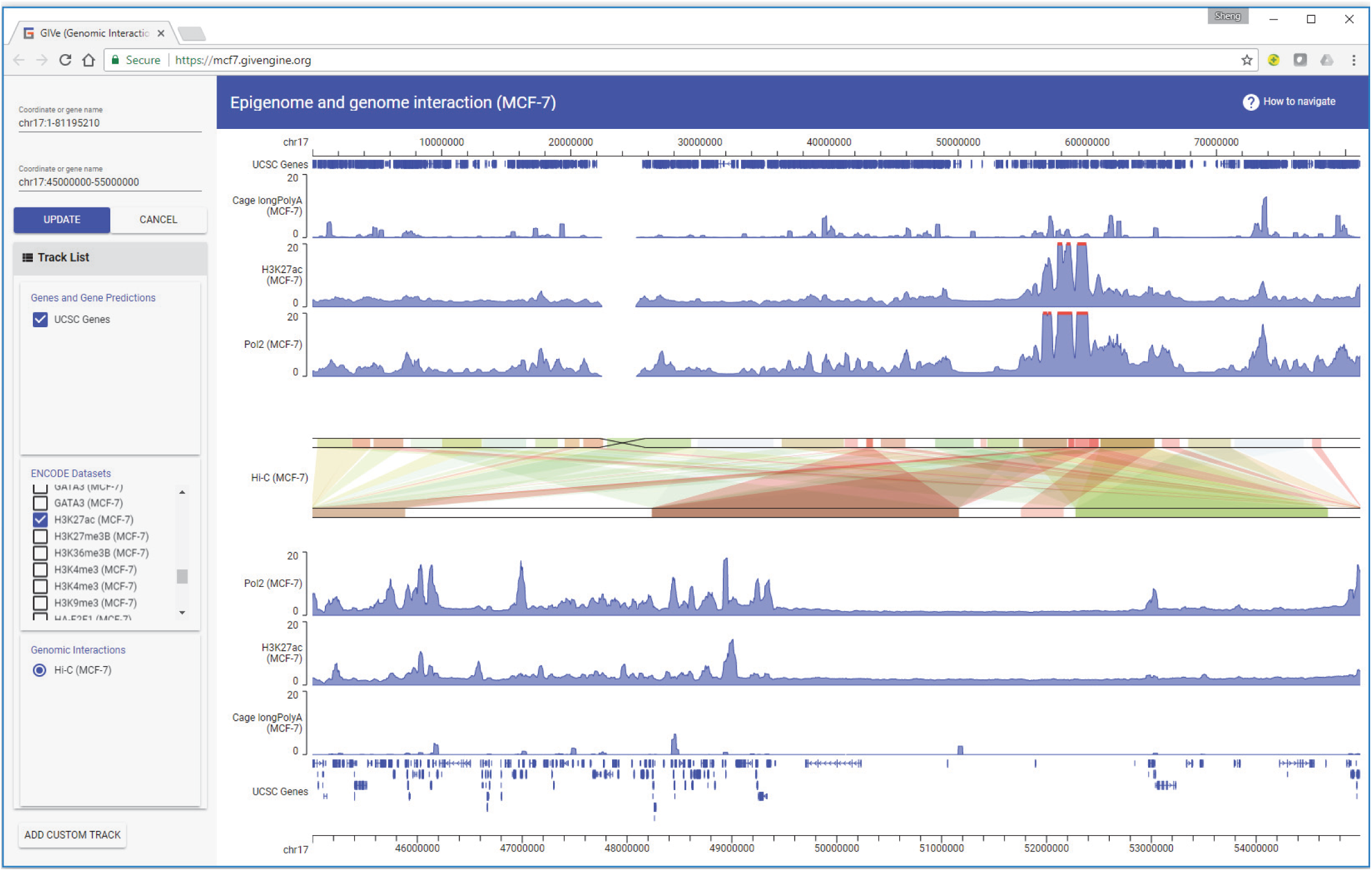
Screenshot of a custom genome browser hosting epigenome and genome interaction datasets. The top genomic coordinate covers the entire chromosome 17 (chr17:1-81195210). The first three data tracks from the top are RNA-seq, H3K27ac ChIP-seq, Pol2 ChIP-seq data in MCF-7 cells, shown corresponding to the top genomic coordinate. The bottom genomic coordinate at the shows chr17:45000000-55000000. The bottom three data tracks are RNA-seq, H3K27ac ChIP-seq, Pol2 ChIP-seq data shown corresponding to the bottom coordinate. The Hi-C interaction data in the center panel shows Hi-C derived links between the genomic regions (top coordinate) to other genomic regions (bottom coordinate). The strengths of the Hi-C derived genomic interactions are plotted in color scale, with red being strongest and green being weakest.

**Figure 2.**
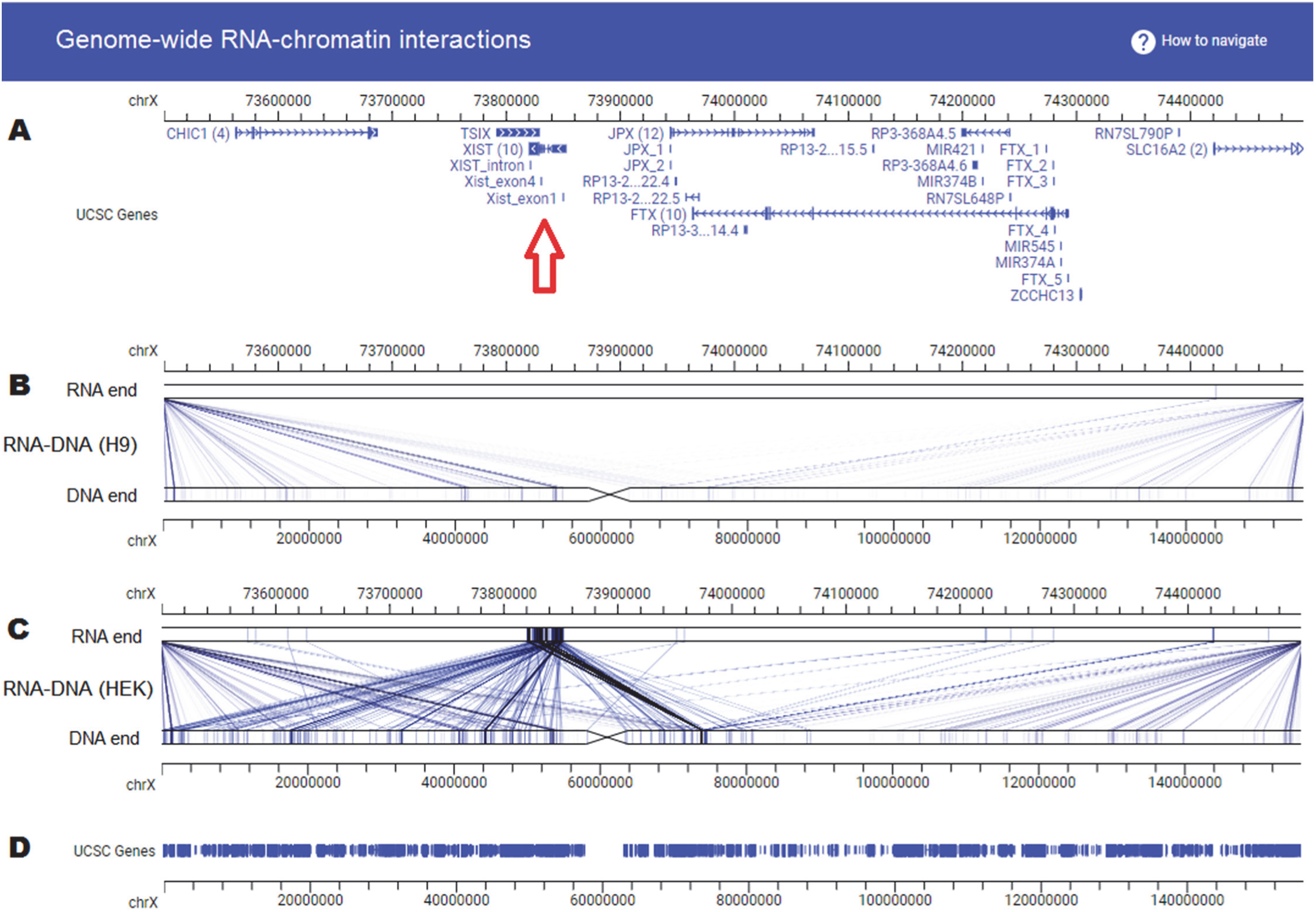
A custom website hosting genome-wide RNA-DNA interaction datasets. Panels from top to bottom are (A) genome coordinates (chrX:73500000-74500000) and Genes, (B) RNA-DNA interaction data in human embryonic (H9) stem cells, with the RNA end (top) and the DNA end (bottom) shown with different resolutions (coordinate bars), (C) RNA-DNA interaction data in human embryonic kidney (HEK) cells, with the RNA end (top) and the DNA end (bottom) shown with different resolutions (coordinate bars), (D) genes and genome coordinates. Red arrow points to the genomic location of the Xist gene, where no RNA was produced in H9 (B) but plenty of RNA was produced and interact with X chromosome is HEK (C). Data were produced by the MARGI technology (Bharat et al., 2017).

### Automatic webpage generation with GIVE HUG (<u>H</u>TML <u>U</u>niversal <u>G</u>enerator)

GIVE data hub and its embedded feature HUG enable automatic generation of interactive visualization webpages for user chosen datasets. GIVE data hub is a web page for browsing the metadata of genomic datasets hosted on public data servers (Figure S2). Inside this web page is a database of metadata, including data type, data description, and the web address of the actual dataset. When a user browses GIVE data hub, the metadata are retrieved and displayed on-demand, ensuring consistency of displayed information with the underlying database. All metadata in GIVE data hub are validated by GIVE development team to ensure correctness of information. Users are welcome to submit metadata of additional datasets hosted on public data servers through an online metadata submission form.

HUG automatically generates HTML webpages for any user chosen datasets. To use HUG, users can click “HTML Generator Mode” in data hub website (Figure S2), select any datasets and click the “Generate” button (Figure S3). A separate window will pop up that summarizes the user chosen datasets and provide the generated HTML code (Figure S4). Like Google map, this data-containing genome visualization HTML code can be copy-pasted into a personal website or saved and shared. Users can interactively change a few display parameters using the top portion of this interactive window and hit “Update code” button, leading to a new HTML code incorporating user-designated visualization parameters (Figure S4). HUG offers the simplest way of generating GIVE-powered genome browser websites.

GIVE data hub and HUG also serve as an interactive tutorial for adding and managing datasets with GIVE-Toolbox and using GIVE’s HTML tags for data visualization, which will be discussed in the next section.

### Managing custom data with GIVE-Toolbox

To add and manage custom data, users should first download and run GIVE’s main executable called GIVE-Docker. GIVE-Docker can be executed on all mainstream operating systems without system specific configuration. When executed, GIVE-Docker automatically sets up a web server and a database system. Also packaged within this executable is a toolbox (GIVE-Toolbox) that automates all database operations into command line commands (Table S1), thus relieving the user from working with a database language. Using the website hosting single-cell transcriptomes (https://singlecell.givengine.org/) as an example, we provide a line-by-line example of building a website hosting custom data. After downloading and running GIVE-Docker, we will issue GIVE-Toolbox provided commands to initialize a reference genome, add gene annotations, and load custom data (Table S2), following by inserting HTML tags to display the data (Last row, Table S2).

Without additional coding, the website is automatically equipped with a few interactive features. These features were enabled by JavaScript codes that were encapsulated within the GIVE’s HTML tags. Visitors to this website can input new genome coordinates (Figure S5A), choose any subset of data tracks to display (Figure S5B), or change genome coordinates by dragging the coordinates to left or right by mouse (Figure S5C) or zoom in or out the genome by scrolling the mouse wheel while the mouse pointer is on top of the genome coordinate area (Figure S5C).

### Double layer display of genome interaction data

GIVE implements a double layer display strategy for visualization of genome interaction data. In this display format, two genomic coordinates are plotted in parallel (central panel of Figure 1, Panels B and C of Figure 2). Interactions between genomic regions are displayed as links of correspondent genomic regions between the top and bottom coordinates. When intensity values are associated with the links, the intensities are displayed in a red (large) to green (small) color scale (central panel, Figure 1). This double layer display strategy has two advantages. First, the top and the bottom coordinates can cover different genomic regions, making it flexible to visualize long range interactions (Figure 1). Users can shift or zoom the top and the bottom coordinates independently, making it easy to visualize for example interactions from the XIST locus (RNA end, Figure 2B-C) to the entire X chromosome (DNA end, Figure 2B-C). This double layer design also makes it intuitive to display asymmetric interactions, for example interactions from RNA (top lanes, Figure 2) to DNA (bottom lanes, Figure 2).

### New data structures for transfer and visualization of genomic data

We developed two data structures for optimal speed in transferring and visualizing genomic data. These data structures and their associated technologies are essential to GIVE. However, all the technologies described in this section are behind the scene. A website developer who uses GIVE does not have to recognize the existence of these data structures.

We will introduce the rationales for developing the new data structures with a use scenario. When a user browses a genomic region, all genome annotation and data tracks within this genomic region should be transferred from the web server to the user’s computer. At this moment, only the data within this genomic region require transfer and display (Figure 3A). Next, the user shifts the genomic region to the left or right. Ideally, the previous data in user’s computer should be re-utilized without transferring again, and only the new data in the additional genomic region should be transferred. After data transfer, the previous data and the new data in user’s computer should be combined (Figure 3B).

**Figure 3.**
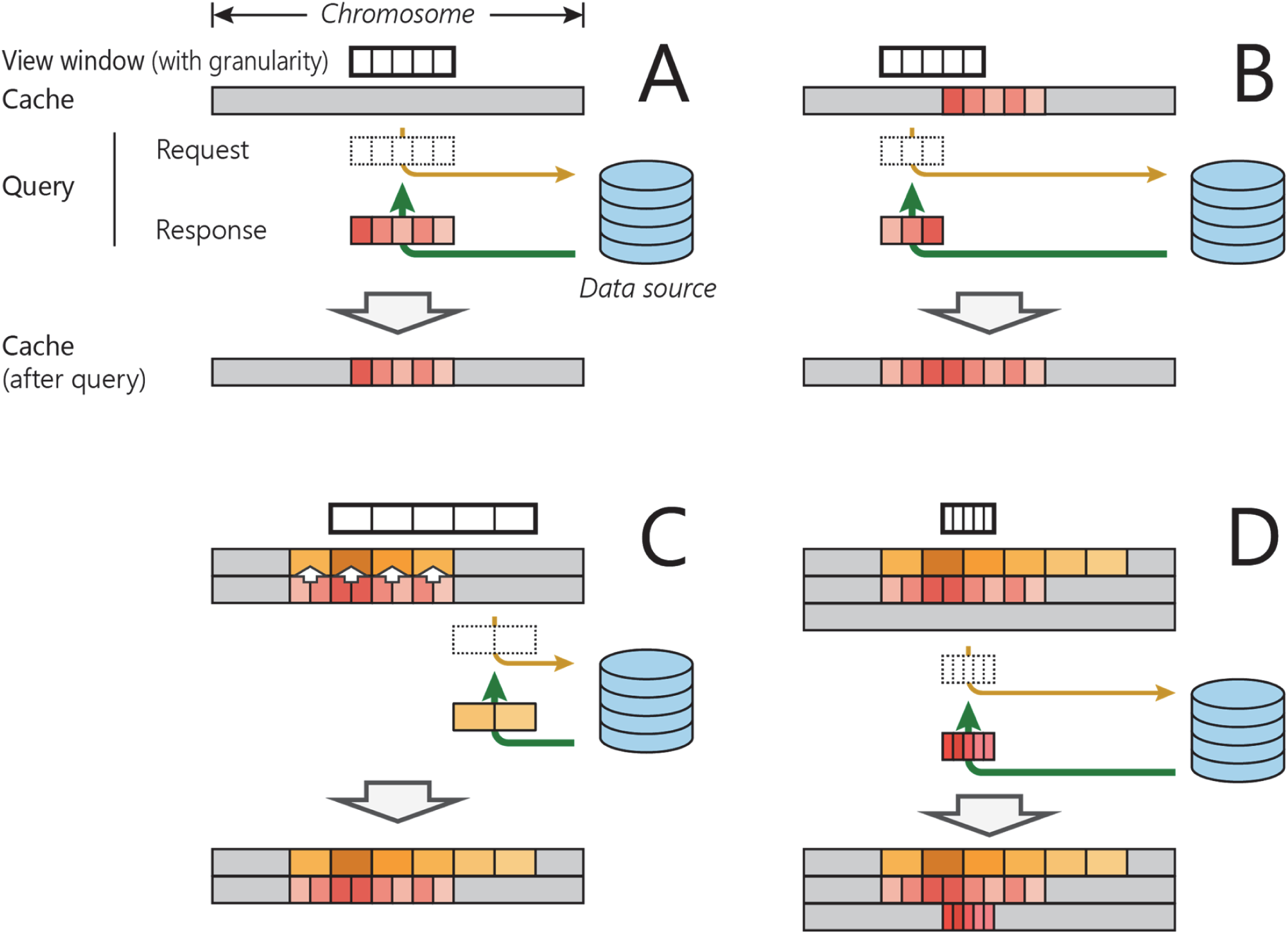
Scenarios for browser use. A) Displaying a segment of the genome. While no data is stored in cache (blank blocks), only those within the queried region needs to be fetched from the server (colored blocks) and is stored in cache for later use; B) Shifting display window. Only the part not in cache needs to be fetched from the server (colored blocks) and merged in cache; C) Zooming out. Existing cache data are used to recalculate new cache at a coarser granularity level, after which non-overlapping data are requested; D) Zooming in. Because no cached data exists at a finer granularity level, all data within the queried region needs to be fetched at that level.

Next, the user zooms out. This action changes the resolution of genome. It is unnecessary to transfer and infeasible to display data at the previous granularity. At this point, the program should adjust the granularity of the already transferred data, and then transfer additional data at the new granularity (Figure 3C). When the user zooms in, the program would adjust to finer granularity, and transfer data at this resolution (Figure 3D). In summary, what is needed is a multiscale data container that can add or remove data from both sides of a genomic window.

To substantiate the above described multiscale data container, we developed two data structures named Oak and Pine. Oak handles sparse data tracks such as genome annotation, gene tracks, peak tracks, and interaction regions (BED, interaction data). Pine deals with dense data track in bigWig format [28]). Once the user changes the viewing area, Oak and Pine automatically adjust to the optimal tree structure for holding the data in the viewing area, which may involve change of data granularity, change of tree depths, adding or merging nodes, and rearrangement node assignment to branches. These operations minimize data transfer over the internet as well as the amount of data loaded in computer memory.

To optimize the use of memory, we developed an algorithm for removing obsolete data from the memory (“withering”). When the data stored in Oak or Pine nodes have not been accessed by the user for a long time, data in these nodes will be dumped and memory is recycled.

## METHODS

### Using HTML tag library

Use of GIVE’s HTML tags do not require any downloading or installation. The simplest way of trying out GIVE’s HTML tags is to use HUG, a graphical interface that will generate an HTML file for user chosen datasets.

Instead of using HUG, a web developer can import the entire GIVE library to a web page by inserting the following two lines (Lines 1, 2).

<script src=“https://www.givengine.org/bower_components/webcomponentsjs/webcomponents-lite.min.js”></script> (Line 1)

<link rel=“import” href=“https://www.givengine.org/components/chart-controller/chart-controller.html“> (Line 2)

To display genomics data, the web developer can use either the <chart-controller> tag or the <chart-area> tag. The <chart-controller> tag will display genomic data as well as genome navigation features such as shifting, zooming (Figure S5C). For example, adding the following line in addition to the two lines above would create a website similar to that in Figure S5 (Line 3).

<chart-controller title-text=“Single-cell RNA-Seq” group-id-list=‘[“genes”, “singleCell”, “customTracks”]’ num-of-subs=“1”></chart-controller> (Line 3)

Here, the title-text attribute sets the title text of a website. The <chart-area> tag will display the track data without metadata controls such as data selection buttons and input box for genomic coordinates, while retaining some interactive capacities including dragging and zooming. This option provides the developer greater flexibility for website design. In addition, the <chart-area> tag is compatible with mobile apps.

### Using GIVE-Toolbox

GIVE-Toolbox is a set of command line tools offered to manage custom data (Table S1). These command line tools automate data related operations and relieve website developers from directly programming with a database language (MySQL). In addition to comprehensive documentation and tutorials (Table S3), executing each tool with –h argument will output usage instruction. GIVE-Toolbox is our recommended option; however, developers can choose to directly work MySQL instead.

### Running GIVE-Docker as a standalone executable

Utilizing Docker’s container technology (https://www.docker.com), we encapsulated GIVE’s codes and all the environmental requirements and database including Apache, MySQL, PHP into a fully packaged executable called GIVE-Docker. This standardized executable can be deployed without system specific configuration to all mainstream operating systems and cloud computing services, including Linux, macOS, Windows 10, AWS, and Azure. This standalone executable does not require system administration or installation of any prerequisite compiler or database, and therefore is the recommended option. Use of GIVE HTML tag library does not require running GIVE-Docker.

Experienced programmers can choose custom installation instead of using GIVE-Docker. A step by step guide of custom installation is provided in GIVE’s online manual.

### Backstage technologies

The following technologies are wrapped inside the GIVE library. Website developers who use GIVE do not have to understand them or even knowing their existence.

### Query

A query is issued when the user views any genomic region (query region). A new query is issued when the user changes the genomic region. A query induces two actions, which are data retrieval and display of data.

### Oak, a data structure

A data structure called Oak is developed to effectively load and transfer a subset of data in BED format. The subset is defined as continuous genomic region within a chromosome. Oak is a type of tree data structures, with nodes defined as follows.

A node is composed of a list of key-value pairs and a set of attributes. A key is a pair of starting and ending genomic coordinates, termed left key and right key, respectively. When populated with data, a node keeps the data for a genomic region defined by the first left key and the last right key. The keys in a node partition the genomic region into non-overlapping sub-regions. A node can be either a branch node or a leaf node. Their differences lie in the values. A branch node is a node where the values are other nodes. A leaf node is a node where each value is a set of two lists of data points (Figure S6). Each data point is a row of a BED file. When populated with data, the first list contains all the rows in the BED file where the start position matches the left key. The second list contains all the rows where the start and the end positions cover (span across) the left key. A value in a leaf node can also be empty. Leaf nodes with empty values are used to mark the genomic regions outside the query region.

### Creating an Oak instance, populating data, and updating Oak

An Oak instance will be created, populated with data, or get updated in response to a query. These actions accomplish data transfer from server to user’s computer. Only the data within the queried region will be transferred. Hereafter we will refer an Oak instance as an Oak.

When the query region is on a new chromosome, an Oak will be created as follows. Every unique start position in the BED file that is contained within the query region is used to create a leaf node. The genomic regions on queried chromosome but outside the query region are inserted as pairs of keys and empty values (placeholders) to the nodes with the nearest keys. The leaf nodes are ordered by their first left keys and sequentially linked by their pointers. A root node is created with all the leaf nodes are its children. This initial tree is fed into a self-balancing algorithm [29, 30] to construct a weight balanced tree, thus finishes the construction of an Oak.

When the query region is on a previously queried chromosome, the query region will be compared with the Oak of that chromosome and the overlapping region will be identified. The data of the overlapping region are already loaded in the Oak and therefore for the purpose of saving time this should not be loaded again. The data in the rest of the query region will be loaded to the Oak. This is done by first creating a leaf node for every additional unique start position, removing the placeholder key-value pairs, and adding new placeholder key-value pairs for the rest of the chromosome. The weight balancing algorithm [29] is invoked again to re-balance this Oak. The weight balancing step prepares the Oak for efficient response to future queries.

### Pine, a data structure

A data structure called Pine is developed to effectively load and transfer a subset of data in bigWig format. The subset is defined as continuous genomic region within a chromosome. Pine can automatically determine the data granularity, which avoids transferring data at a higher than necessary resolution. The resolution of displayed data is limited by the number of pixels on the screen. Pine instances are always constructed to the appropriate depth and match the limit of the resolution.

A node consists of a list of key-value pairs and a set of attributes. The attributes are the same as that of Oak nodes, except for having one additional attribute, called data summary. The data summary includes the following metrics for this node (the genomic region defined by the first left key and the last right key of this node): the number of bases, sum of values (summing over every base), sum of squares of the values, maximum value, and minimum value. A key is a pair of starting and ending genomic coordinates, termed left key and right key, respectively. The keys in a node partition the genomic region into non-overlapping sub-regions. A node can be either a branch node or a leaf node. Their differences lie in the values. A branch node is a node where the values are other nodes (Figure S7A). A leaf node is a node where each value is a list of data points (Figure S7B). Each data point is a row (binary format) of a bigWig file.

A node in Pine can have an empty key-value list and an empty data summary, and in this case we call it a placeholder node.

### Creating a Pine instance, populating data, and updating Pine

A Pine is created when a query to a new chromosome is issued. A Pine is created with the following steps. First, the depth of the Pine tree is calculated as:

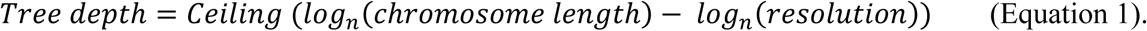

The limit of the resolution (length of genomic region per pixel) is the total length of the queried genomic region (viewing area) divided by the number of horizontal pixels, namely the width of the SVG element in JavaScript.

Next, a root node is created with keys covering the entire chromosome, where the query region is contained within. Until reaching the calculated depth, for any node that overlaps with the query region, create a fixed number (n, n=20 in the current release) of child nodes by equal partitioning its genomic region. If any of the created child node does not overlap with the query region, use a placeholder node. For each node, point the pointer to the “right hand” node at the same depth. Thus, a Pine is created. This Pine has not loaded with actual data.

To load data, every leaf node issues a request to retrieve the summary data of its covered region (between the first left key and the last right key), which will be responded by a PHP function wrapped within GIVE. This function returns summary data between the input coordinates from the bigWig file. After filling the summary data for all nodes at the deepest level, all parent nodes will be filled, where the summary data are calculated from the summary data of their child nodes. This process continues until reaching the root node.

A Pine will be updated when a new query partially overlaps with a previous query. In this case, the new depth (d2) is calculated using Equation 1. This depth (d2) reflects the new data granularity. If d2 is greater than the previous depth, extend the Pine by adding placeholder nodes until d2 is reached. From root to depth d2-1, if any placeholder node overlaps with the query region, partition it by creating n child nodes. If any of the newly created child node does not overlap with the query region, use a placeholder node. For any newly created node, point the pointer to the “right hand” node at the same depth. At this step, the Pine structure is updated into proper depth. Finally, at depth d2, retrieve summary data for every non-placeholder node that has not had summary data. Update the summary data of their parent nodes until reaching the root. In this way, only the new data within the query region that had not been transferred before will get transferred.

### Memory management

We developed a memory management algorithm called “withering”. Every time a query is issued, this algorithm is invoked to dump the obsolete data, which are not used in the previous 10 queries. “Withering” works as follows: all nodes are added with a new integer attribute called ‘life span’. When a node is created, its life span is set to 10. Every time a query is issued, all nodes overlapping with the query region as well as all their ancestral nodes get their life span reset to 10. The other nodes that do not overlap with the query region get their life span reduced by 1. All the nodes with life span equals 0 are replaced by placeholder nodes.

## DISCUSSION

The GIVE library is designed to reduce the need for specialized knowledge and programming time for building web-based genome browsers. GIVE is open source software. The open source nature allows the community at large to contribute to enhancing GIVE. The name GIVE (Genome Interaction Visualization Engine) was given when this project started with a smaller goal. Although it has grown into a more general-purpose library, we have decided to keep the acronym.

An important technical consideration is efficient data transfer between the server and user’s computers. This is because users typically wish to get an instant response when browsing data. To this end, we developed several technologies to optimize the speed of data transfer. The central idea is in three folds, including 1) only transferring the data in the query region, 2) minimizing repeated data transfer by reusing previously transferred data, and 3) only transferring data at the necessary resolution. To implement these ideas, we developed two new approaches to index the genome, and formalized these approaches with two new data structures, named Oak and Pine.

The Oak and Pine are indexing systems for sparse data (BED) and dense data (bigWig), respectively. BED data typically store genomic segments that have variable lengths. Given this particular feature, we did not index the genome base-by-base but rather developed a new strategy (Oak) to index variable-size segments. The bigWig files contain base-by-base data, which for a large genomic region can become too slow for web browsing. We therefore designed the Pine data structure that can automatically assess and adjust data granularity, which exponentially cut down unnecessary data transfer.

## ADDITIONAL INFORMATION

GIVE website is at http://www.givengine.org, which provides samples websites, tutorial, manual, and GIVE executable. A mirror website is at: https://sites.google.com/view/givengine. Source codes are available at GitHub (https://github.com/Zhong-Lab-UCSD/Genomic-Interactive-Visualization-Engine) and at Zenodo (DOI: 10.5281/zenodo.1134907). A total of 9 demos and tutorials with real codes and complete instructions are provided GIVE manual (Table S3) (https://github.com/Zhong-Lab-UCSD/Genomic-Interactive-Visualization-Engine/tree/master/tutorials).

## ACKNOWLEDGEMENT

We thank the open source community especially Adel Qalieh, Yuan Liu and authors of the Polymer programming library [31]. This work is funded by NIH U01CA200147 and DP1HD087990.

## COMPETING INTREST

Sheng Zhong is a cofounder and a board member of Genemo Inc., which however does not do business related to work described in this paper.

