## Supplementary Materials for "GIVE: toward portable genome browsers for personal websites"

### SUPPLEMENTARY FIGURES

Figure S1. Screenshot of a website hosting ENCODE datasets.

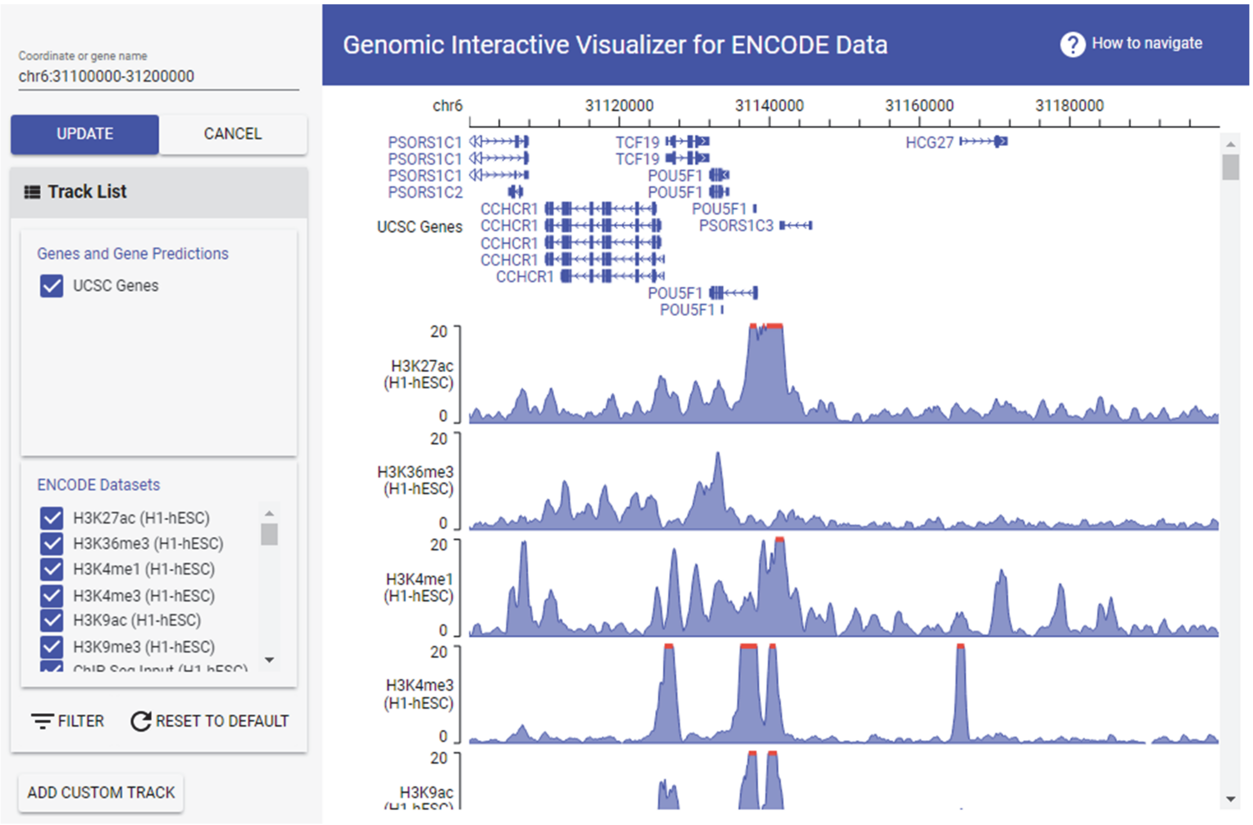

Figure S2. Screenshots of GIVE data hub. Clicking on "HTML generator mode" on top right corner would activate GIVE HUG.

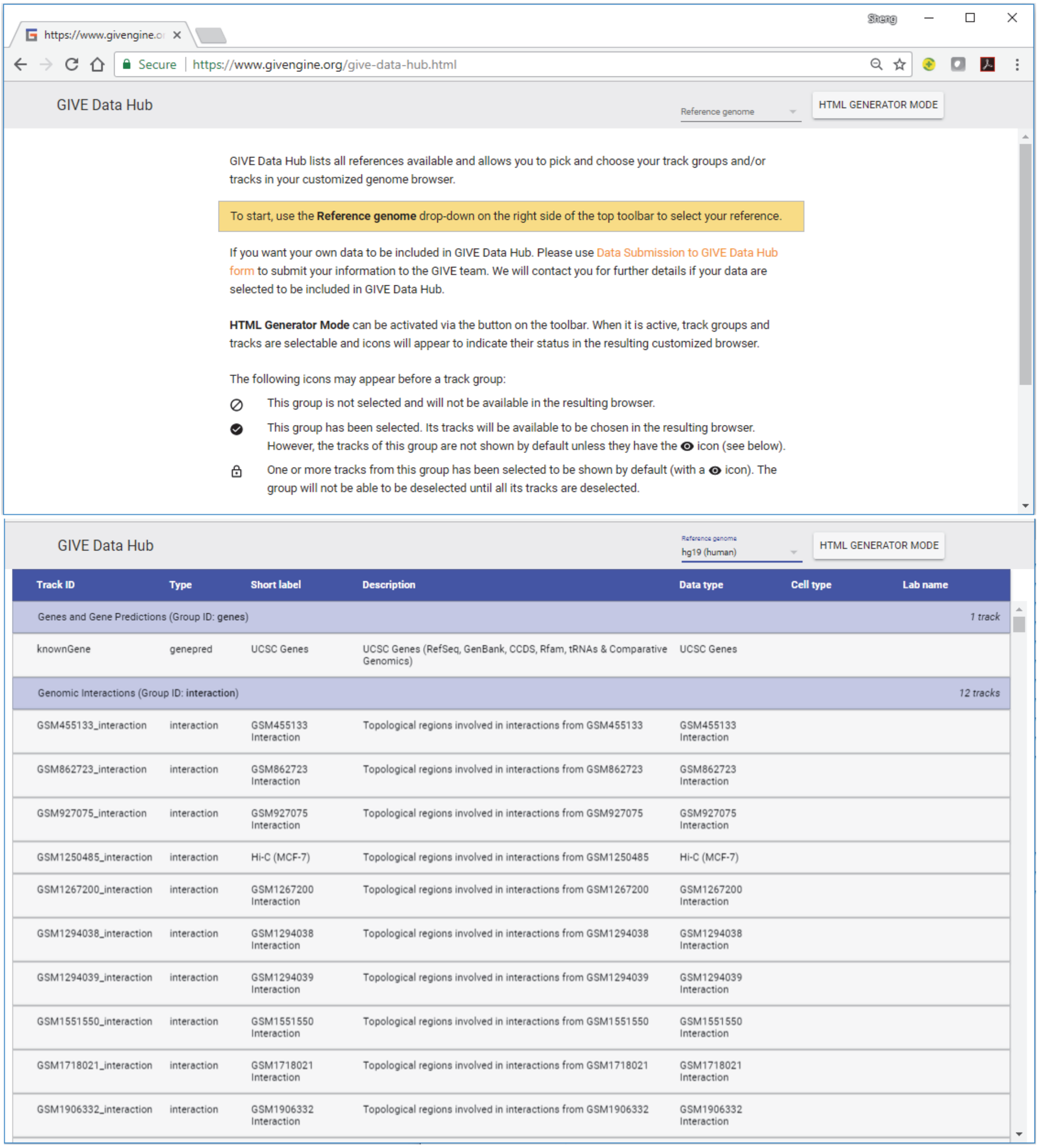

Figure S3. Selection of datasets in GIVE data hub. Clicking on a dataset will get it selected (orange) and clicking again will unselect it (yellow). Clicking on "Generate" button (upper right corner) will produce a HTML file for displaying the selected datasets.

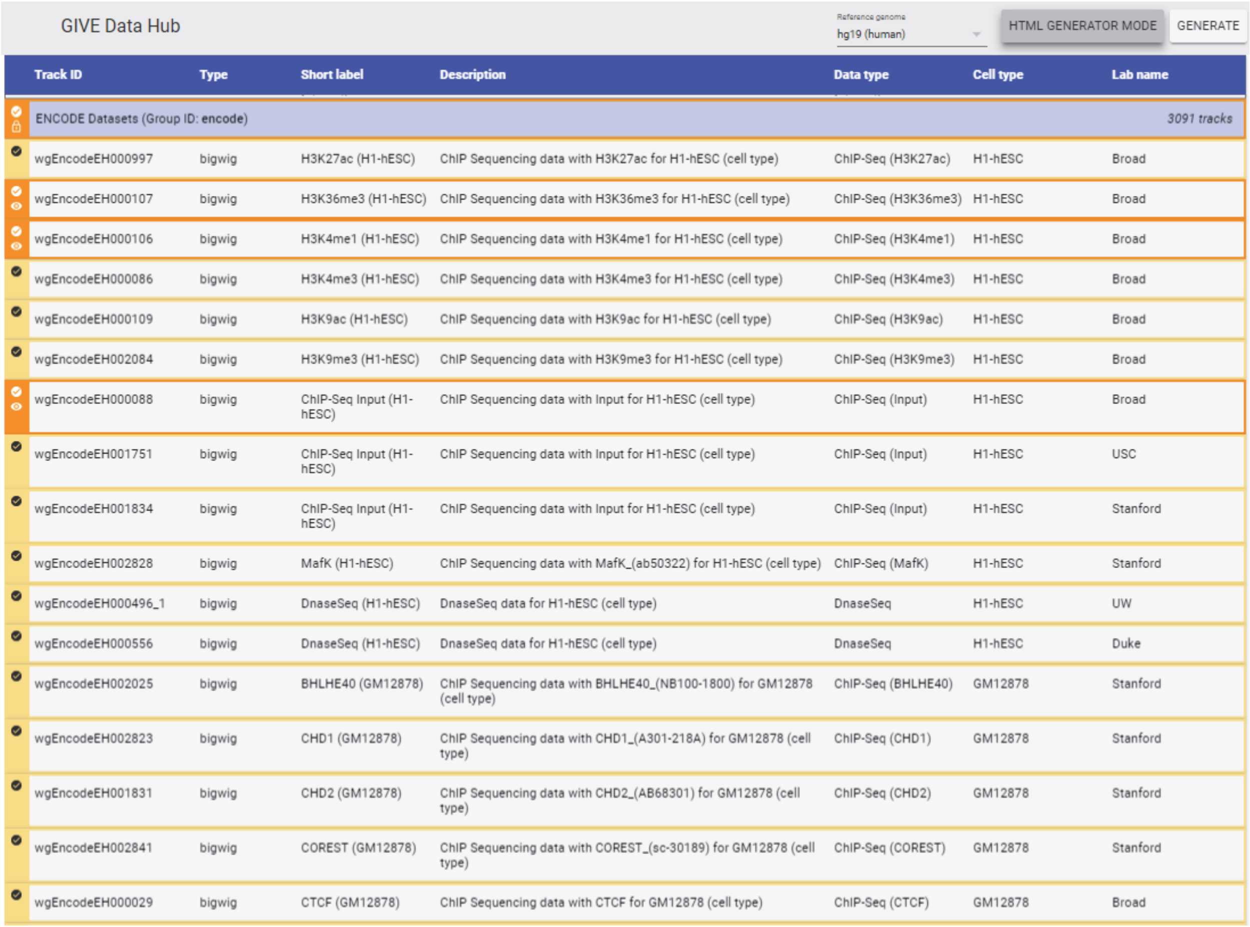
GIVE Data Hub — Reference genome: hg19 (human) — HTML GENERATOR MODE — GENERATE

| Track ID | Type | Short label | Description | Data type | Cell type | Lab name |
|---|---|---|---|---|---|---|
| ENCODE Datasets (Group ID: encode) | | | | | | 3091 tracks |
| wgEncodeEH000997 | bigwig | H3K27ac (H1-hESC) | ChIP Sequencing data with H3K27ac for H1-hESC (cell type) | ChIP-Seq (H3K27ac) | H1-hESC | Broad |
| wgEncodeEH000107 | bigwig | H3K36me3 (H1-hESC) | ChIP Sequencing data with H3K36me3 for H1-hESC (cell type) | ChIP-Seq (H3K36me3) | H1-hESC | Broad |
| wgEncodeEH000106 | bigwig | H3K4me1 (H1-hESC) | ChIP Sequencing data with H3K4me1 for H1-hESC (cell type) | ChIP-Seq (H3K4me1) | H1-hESC | Broad |
| wgEncodeEH000086 | bigwig | H3K4me3 (H1-hESC) | ChIP Sequencing data with H3K4me3 for H1-hESC (cell type) | ChIP-Seq (H3K4me3) | H1-hESC | Broad |
| wgEncodeEH000109 | bigwig | H3K9ac (H1-hESC) | ChIP Sequencing data with H3K9ac for H1-hESC (cell type) | ChIP-Seq (H3K9ac) | H1-hESC | Broad |
| wgEncodeEH002084 | bigwig | H3K9me3 (H1-hESC) | ChIP Sequencing data with H3K9me3 for H1-hESC (cell type) | ChIP-Seq (H3K9me3) | H1-hESC | Broad |
| wgEncodeEH000088 | bigwig | ChIP-Seq Input (H1-hESC) | ChIP Sequencing data with Input for H1-hESC (cell type) | ChIP-Seq (Input) | H1-hESC | Broad |
| wgEncodeEH001751 | bigwig | ChIP-Seq Input (H1-hESC) | ChIP Sequencing data with Input for H1-hESC (cell type) | ChIP-Seq (Input) | H1-hESC | USC |
| wgEncodeEH001834 | bigwig | ChIP-Seq Input (H1-hESC) | ChIP Sequencing data with Input for H1-hESC (cell type) | ChIP-Seq (Input) | H1-hESC | Stanford |
| wgEncodeEH002828 | bigwig | MafK (H1-hESC) | ChIP Sequencing data with MafK_(ab50322) for H1-hESC (cell type) | ChIP-Seq (MafK) | H1-hESC | Stanford |
| wgEncodeEH000496_1 | bigwig | DnaseSeq (H1-hESC) | DnaseSeq data for H1-hESC (cell type) | DnaseSeq | H1-hESC | UW |
| wgEncodeEH000556 | bigwig | DnaseSeq (H1-hESC) | DnaseSeq data for H1-hESC (cell type) | DnaseSeq | H1-hESC | Duke |
| wgEncodeEH002025 | bigwig | BHLHE40 (GM12878) | ChIP Sequencing data with BHLHE40_(NB100-1800) for GM12878 (cell type) | ChIP-Seq (BHLHE40) | GM12878 | Stanford |
| wgEncodeEH002823 | bigwig | CHD1 (GM12878) | ChIP Sequencing data with CHD1_(A301-218A) for GM12878 (cell type) | ChIP-Seq (CHD1) | GM12878 | Stanford |
| wgEncodeEH001831 | bigwig | CHD2 (GM12878) | ChIP Sequencing data with CHD2_(AB68301) for GM12878 (cell type) | ChIP-Seq (CHD2) | GM12878 | Stanford |
| wgEncodeEH002841 | bigwig | COREST (GM12878) | ChIP Sequencing data with COREST_(sc-30189) for GM12878 (cell type) | ChIP-Seq (COREST) | GM12878 | Stanford |
| wgEncodeEH000029 | bigwig | CTCF (GM12878) | ChIP Sequencing data with CTCF for GM12878 (cell type) | ChIP-Seq (CTCF) | GM12878 | Broad |

Figure S4. HUG generated HTML code. Top row summarizes user selected datasets, including reference genome, data group and short descriptions of selected datasets. The generated HTML code is shown in lower half of this window. Users can use the interactive features in the middle portion of this window to modify display parameters. Clicking “Update code” button would refresh the lower half with a regenerated code incorporating the new parameters.

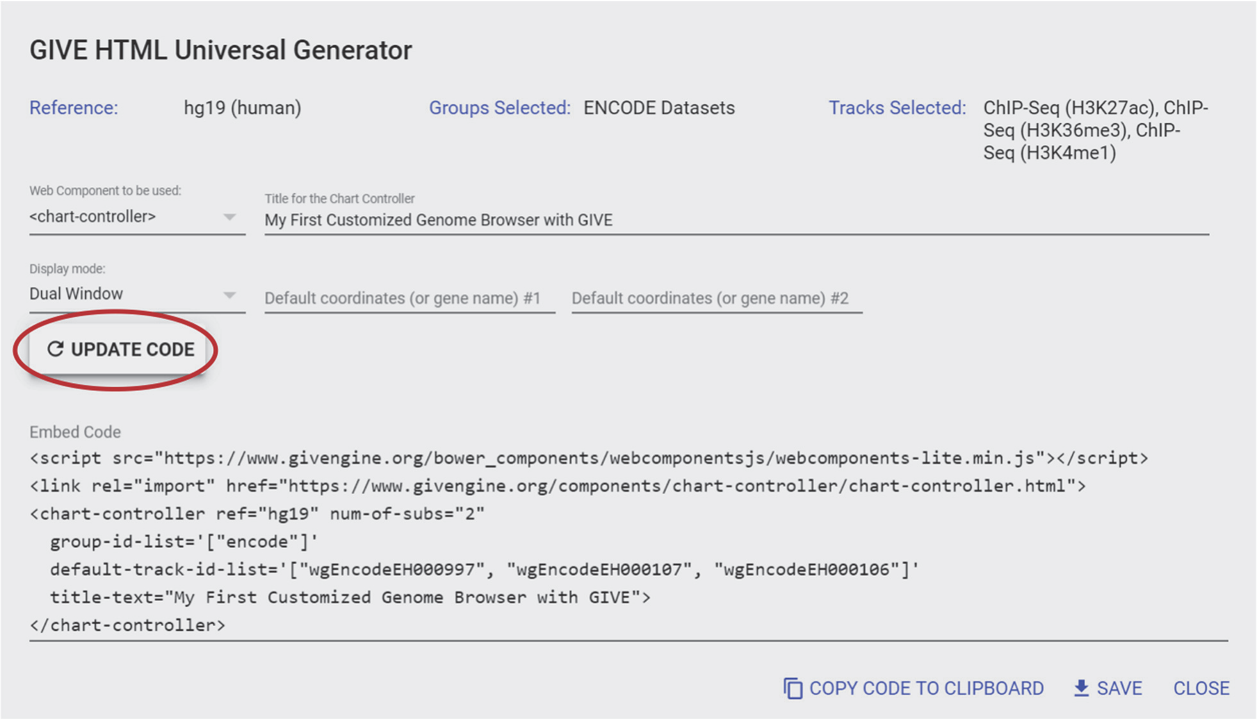

Figure S5. Screenshot of a custom genome browser. (A) Text box and "update" bottom for changing genomic coordinates. (B) Check boxes for choices of datasets for retrieval and display. (C) Genome coordinates and genes. Dragging with mouse would shift genome coordinates. Rolling the mouse wheel would zoom in or out.

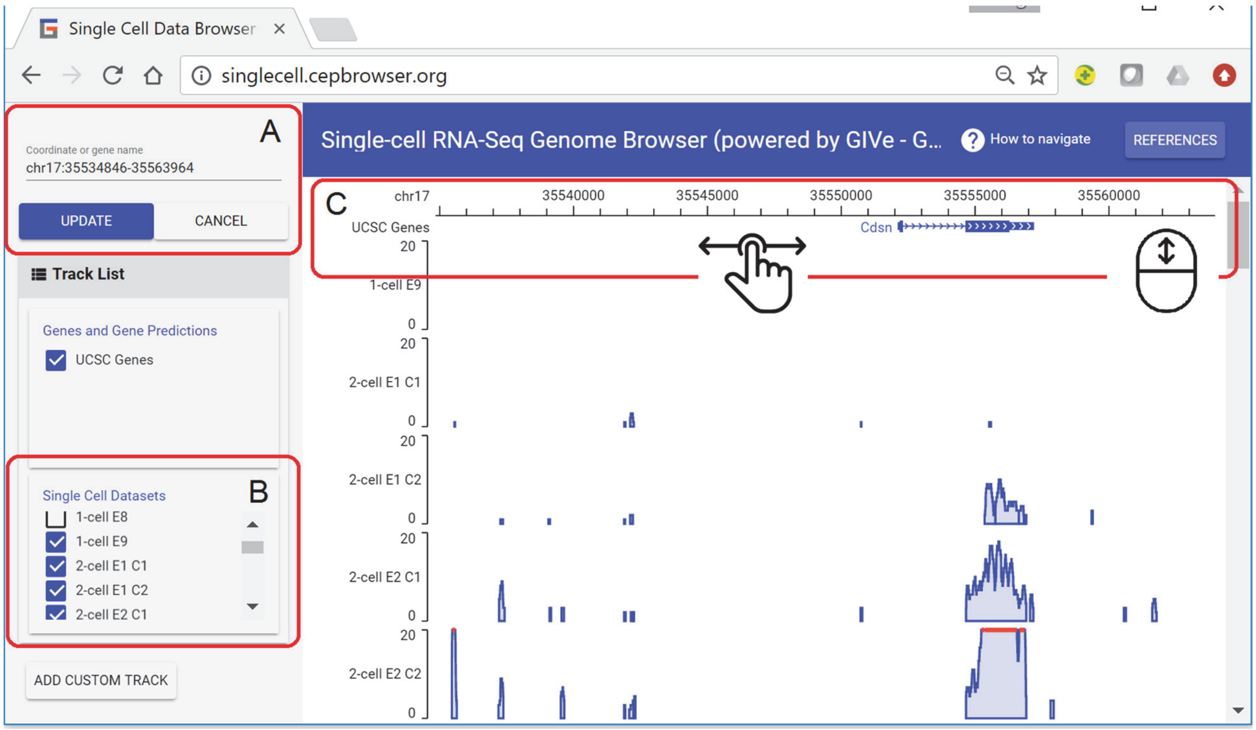

Figure S6. Oak data structure and operations.

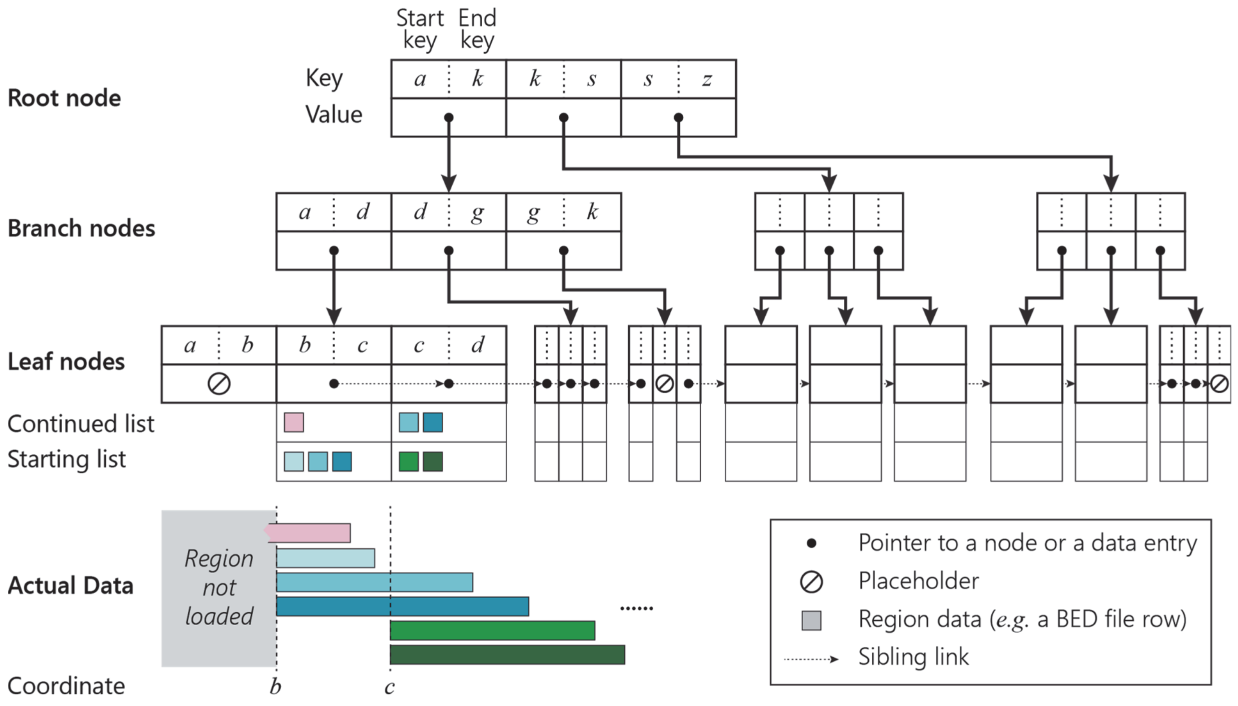

Figure S7. Pine data structure and operations.

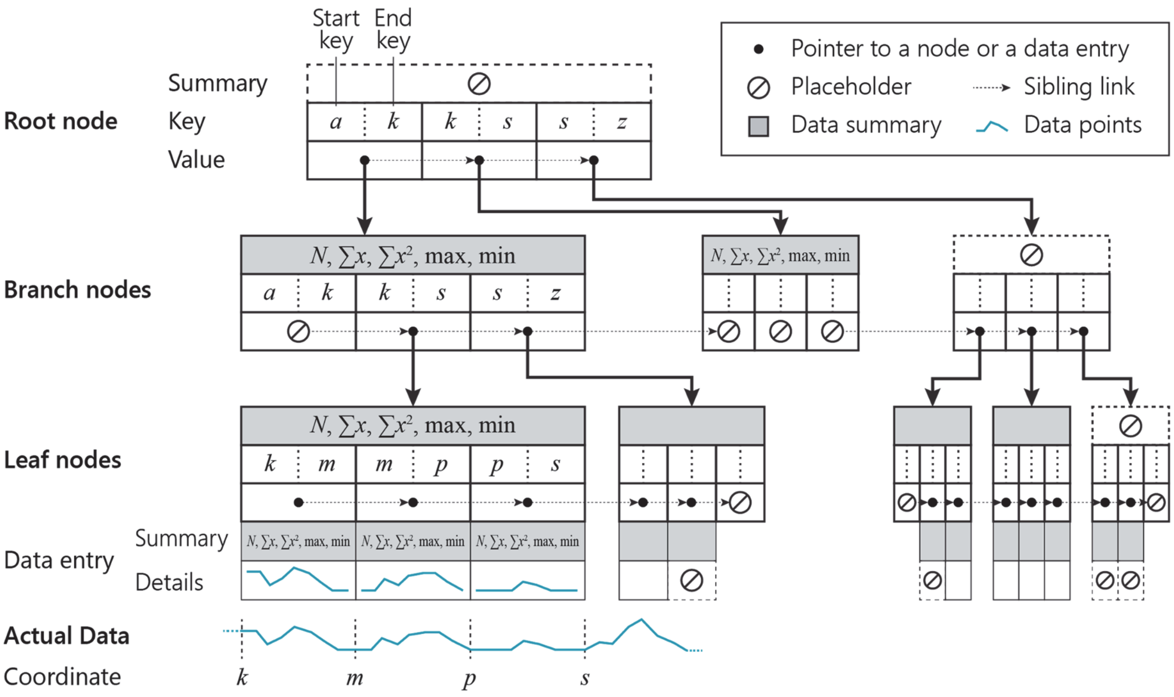

### SUPPLEMENTARY TABLES

Table S1. Summary of GIVE Toolbox. GIVE Toolbox is a set of command line commands (Command) that automated databased operations (Operations).

| Goal | Command | Operations |
|---|---|---|
| **Initialize a reference genome** | initial_ref.sh | Create a database of the reference genome |
| | | Create a table for the chromosome sizes |
| | | Import data to the chromosome sizes table |
| | | Register the reference genome database |
| | | Create a table for track group registration |
| | | Create a table for track registration |
| **Create a data track** | add_trackGroup.sh | Register the track group that the track belongs to |
| | (choose based on data type)<br>add_geneAnnot.sh<br>add_track_bed.sh<br>add_track_bigWig.sh<br>add_track_interaction.sh | Create a table for the data track |
| | | Import data to the data track table |
| | | Register and add annotation (metadata) of the data track |
| **List existing data** | list_tracks.sh | Check registration tables of reference genome, track groups and data tracks one by one |
| **Remove data** | remove_data.sh | Drop the data track table |
| | | Delete the info in related registration table |

Table S2. Related to Figure 1. Line-by-line commands and codes for creating a genome browser loaded with custom data. The first three steps (Step column) were automated as command line commands (Command/HTML tag) in GIVE-Toolbox (Component). The final step utilizes an HTML tag provided in the HTML tag library (Component).

| Step | Line by line command or code | Component | Command/HTML tag |
|---|---|---|---|
| **Initiate a reference genome** | Create data files for the reference genome | GIVE-Toolbox | initial_ref.sh |
| **Add gene annotations** | Create a track group for gene annotations | | add_trackGroup.sh |
| | Add gene annotation file | | add_geneAnnot.sh |
| **Load custom data** | Create a track group for custom data tracks | | add_trackGroup.sh |
| | Load a custom data file | | add_track_bigWig.sh |
| | Repeat the last step to add other data files | | add_track_bigWig.sh |
| **Display data** | Insert an HTML tag to display the data | HTML tag library | <chart-controller> |

Table S3. Templates with real codes and complete instructions. All these demos are accessible by the hyperlinks in this table and available in “GIVE Tutorial” from GIVE’s homepage (www.givengine.org).

| Number | Title |
|---|---|
| 1 | Start from a 2-minute example |
| 2 | Build a genome browser with GIVE data hub |
| 3 | Local deployment of GIVE with GIVE-Docker |
| 4 | Use GIVE-Toolbox to manage custom data |
| 5 | Demo of long-range promoter-enhancer interactions with capture Hi-C |
| 6 | Demo with ChIA-PET derived chromatin interactions |
| 7 | Demo with superimposed wiggle and genomic interaction tracks |
